# Gastrointestinal ulceration in calves presented to a central Iowa veterinary referral facility: An underappreciated morbidity?

**DOI:** 10.1101/2020.12.17.423269

**Authors:** Allison Mosichuk, Joe Smith, Dane Tatarniuk, Amanda Kreuder

## Abstract

The objective of this retrospective investigation is to identify the incidence of gastrointestinal ulceration as a co-morbidity in calves presenting to a referral veterinary hospital. Approximately 24% of calves presented to the hospital that died or were euthanized had evidence of gastrointestinal ulceration. Previous administration of an NSAID was significantly associated with the presence of ulcers, whereas antibiotic administration, age at presentation, gender, or breed were not. Clinicians and producers should consider the risk of ulceration in calves treated with NSAIDs, in light of antiulcer therapies.

## Introduction

Gastrointestinal ulceration is a common problem across species, especially horse, pigs, and ruminants. While the common causes include type of diet, diet changes, stress, medications, and infectious agents, there are numerous potential causes. Depending on the severity of the ulceration, animals may be asymptomatic or have serious consequences including maldigestion, bleeding, gastrointestinal perforation, and sudden death.(1) Gastrointestinal ulceration can also lead to decreased weight gain, production loss, and decreased production quality, which can have a significant impact on food animal species. Diagnosis, particularly in ruminants, can be challenging, especially if no clinical signs are present. Ulceration is often undiagnosed and discovered during post-mortem examination.(2)

Treatment for gastrointestinal ulceration is often not done due to the challenges of diagnosis. Direct visualization via endoscopy is challenging and expensive. Other testing strategies, such as fecal occult blood testing, lack sensitivity. If clinical signs are present and ulcers are suspected, ruminants can be treated with H2-receptor antagonists or proton pump inhibitors, which help to increase the pH of the abomasum. The goal of this study was to identify risk factors at necropsy to guide clinicians and producers when deciding if a calf is at risk for gastrointestinal ulceration.

## Materials and Methods

Medical records of the Food Animal and Camelid Hospital of the Iowa State University Lloyd Veterinary Medical Center (ISU LVMC), were generated for all calves under 90 days of age that were either died or euthanized over a period of time from January 1^st^, 2015 through May 1^st^, 2020. These records were analyzed for demographic information, antibiotic and anti-inflammatory administration, and diagnosis. Necropsy reports were also analyzed for pathology of the gastrointestinal tract, lungs, liver, kidneys, joints, brain, and spinal cord. The presence of any form of gastrointestinal ulcers was recorded.

Calves were divided into 2 groups: those with evidence of gastrointestinal ulceration and those without. These groups were then statistically compared to determine if any of the following led to an increased incidence of abomasal ulceration: NSAID or antibiotic administration, age (< 30 days of age), breed (dairy or beef), or gender (male or female). A Fisher exact test was used for comparison (<u>Easy Fisher Exact Test Calculator</u>. (socscistatistics.com), and a *P* < 0.05 was considered statistically significant.

## Results and Discussion

Of the 130 calves that were admitted to ISU LVMC and were sent to necropsy, thirty-one had evidence of gastrointestinal ulceration and ninety-nine did not.

The average age of calves with gastrointestinal ulceration is 15 days +/− 21 days and the average weight is 47.6kg +/− 14.3kg. Twenty-seven of the calves are beef breeds (mixed breed beef (*n* = 15), red angus (*n* = 4), Aberdeen angus (*n* = 3), Hereford (*n* = 2), Simmental (*n* = 1), Shorthorn (*n* = 1), and Maine Anjou (*n* =1)) and four of the calves are dairy breeds (Holstein (*n* = 4)). Twenty-one of the calves are male and ten of the calves are female. Nineteen of these calves were euthanized and twelve of them died naturally as a progression of their disease.

The average age of calves without gastrointestinal ulceration is 17 days +/− 20 days and the average weight is 48.4kg +/− 31.8kg. Sixty-eight of these calves are beef breeds (mixed breed beef (*n* = 40), Aberdeen angus (*n* = 15), Hereford (*n* = 4), Maine Anjou (*n* = 3), Charolais (*n* = 2), Chianina (*n* =1), Gelbvieh (*n* = 1), red angus (*n* = 1) and White Park (*n* = 1)) and thirty-one of the calves are dairy breeds (Holstein (*n* = 15), Simmental (*n* = 9), Shorthorn (*n* = 5), Brown Swiss (*n* = 1), and Jersey (*n* = 1)). Forty-nine of these calves are male and fifty of the calves are female. Sixty-two of these calves were euthanized and thirty-seven of them died naturally as a progression of their disease.

Many of the calves were treated with antibiotics (*n* = 79) or anti-inflammatories (*n* = 57) prior to necropsyfluino. Of the group with evidence of gastrointestinal ulceration 19/31 were treated with an NSAID. Of the group without evidence of gastrointestinal ulceration 38/99 were treated with an NSAID. NSAIDs used included flunixin meglumine (ulceration group (*n* = 10) and no ulceration group (*n* = 29)), meloxicam (ulceration group (*n* = 5) and no ulceration group (*n* = 7)), both (ulceration group (*n* = 4) and no ulceration group (*n* = 1)) or unknown (no ulceration group (*n* = 1)). Calves administered an NSAID were more likely to have evidence of gastrointestinal ulceration on necropsy (*P* = 0.0372).

Of the group with evidence of gastrointestinal ulceration 23/31 were treated with an antibiotic. Antibiotics used included ampicillin (*n* = 9), florfenicol (*n* = 6), potassium penicillin (*n* = 4), ceftiofur sodium (*n* = 2), penicillin (*n* = 2), sulfamethazine (*n* = 2), tulathromycin (*n* = 1), tilmicosin (*n* = 1), amikacin (*n* = 1), and unknown (*n* = 1). Of the group without evidence of gastrointestinal ulceration 56/99 were treated with an antibiotic. Antibiotics used included florfenicol (*n* = 19), ampicillin (*n* = 17), tulathromycin (*n* = 13), ceftiofur hydrochloride (*n* = 9), potassium penicillin (*n* = 7), enrofloxacin (*n* = 4) penicillin (*n* = 3), sulfamethazine (*n* = 2), amikacin (*n* = 2), ceftiofur (*n* = 2) and gentamicin (*n* = 1). No significance was noted from the administration of an antibiotic on the likelihood to have evidence of gastrointestinal ulceration on necropsy (*P* = 0.0941).

When age was evaluated it was noted that of the group with evidence of gastrointestinal ulceration 24/31 were less than 30 days of age. It was noted that in the group with no evidence of ulceration 73/99 were less than 30 days of age. Age was not significant for likelihood of evidence of gastrointestinal ulceration on necropsy (*P* = 0.8147).

When breed was evaluated for evidence of gastrointestinal ulceration 4/31 with ulceration were dairy breeds. Of the group without evidence of gastrointestinal ulceration 17/99 were dairy breeds. Breed was not found to be a significant finding for the likelihood to have evidence of gastrointestinal ulceration on necropsy (*P* = 0.7809).

When gender was evaluated with respect to evidence of gastrointestinal ulceration 19/31 with ulceration were male. Of the group without evidence of gastrointestinal ulceration 49/99 were male. No significant relationships were noted for likelihood to have evidence of gastrointestinal ulceration on necropsy based on gender (*P* = 0.0988). Percentages and P values for all comparisons are listed in **Table 1**.

**Table 1:** Percentages and P values of different clinical variables in calves with evidence of gastrointestinal ulceration on necropsy. P < 0.05 considered statistically significant (bolded).

| Variable | Calves with Ulcers | Calves without Ulcers | P value |
| --- | --- | --- | --- |
| NSAID Administration | 19/31 (61.3%) | 38/99 (38.4%) | **0.0372** |
| Antibiotic Administration | 23/31 (74.2%) | 56/99 (56.6%) | 0.0941 |
| Age (< 30 days) | 24/31 (77.4%) | 73/99 (73.7%) | 0.8147 |
| Breed (dairy) | 4/31 (19.4%) | 17/99 (17.2%) | 0.7809 |
| Gender (male) | 21/31 (67.7%) | 49/99 49.5%) | 0.0988 |

## Conclusions

Of the examined variables in this investigation the only significant relationship between the development of gastrointestinal ulceration was NSAID administration. While NSAIDs reduce the production of prostaglandins in order to decrease inflammation associated with pain, this can also lead to gastrointestinal ulceration and erosion. This consistent in this study where calves with evidence of gastrointestinal ulceration has a significantly higher NSAID use compared to the group with no evidence of ulceration.

This study was limited by a small population of animals presenting to a single referral hospital, as well as economic factors involving individual patient care were not consistent amongst study animals.

Future directions of this work include evaluation of specific therapies (antibiotics, anti-inflammatories, and other supportive care) on development of gastrointestinal ulcers in calves.

Based on this information, clinicians should be cognizant of the increased risk of ulcer development in calves less than 90 days of age presented to a veterinary teaching hospital that were administered an NSAID. Judicious use, clinical monitoring, and gastroprotectant therapy may be indicated for calves of this demographic.

## Notes

### Competing Interest Statement

The authors have declared no competing interest.

